# Quantitative RNA-seq meta analysis of alternative exon usage in *C. elegans*

**DOI:** 10.1101/134718

**Authors:** Nicolas Tourasse, Jonathan R. M. Millet, Denis Dupuy

## Abstract

Almost twenty years after the completion of the *C. elegans* genome sequence, gene structure annotation is still an ongoing process with new evidence for gene variants still being regularly uncovered by additional in-depth transcriptome studies. While alternative splice forms can allow a single gene to encode several functional isoforms the question of how much spurious splicing is tolerated is still heavily debated.

Here we gathered a compendium of 1,682 publicly available *C. elegans* RNA-seq datasets to increase the dynamic range of detection of RNA isoforms and obtained robust measurements of the relative abundance of each splicing event. While most of the splicing reads come from reproducibly detected splicing events, a large fraction of purported junctions are only supported by a very low number of reads. We devised an automated curation method that takes into account the expression level of each gene to discriminate robust splicing events from potential biological noise. We found that rarely used splice sites disproportionately come from highly expressed genes and are significantly less conserved in other nematode genomes than splice sites with a higher usage frequency.

Our increased detection power confirmed trans-splicing for at least 84% of *C. elegans* protein coding genes. The genes for which trans-splicing was not observed are overwhelmingly low expression genes, suggesting that the mechanism is pervasive but not full captured by organism-wide RNA-Seq.

We generated annotated gene models including quantitative exon usage information for the entire *C. elegans* genome. This allows users to visualize at a glance the relative expression of each isoform for their gene of interest.

## INTRODUCTION

In multicellular organisms, cell differentiation is driven by proteomic diversity. Each cell-type and tissue is defined initially by selective expression of gene subsets from the shared genome. Additionally, it is possible for an expressed gene to be subjected to alternative splicing, such that a different subset of exons can be retained or excluded in the final protein-coding mRNAs. Alternative splicing thus allows a single gene to encode several protein variants, called isoforms, with altered stability, localization, specificity or activity (Kelemen et al., 2013).

The importance of alternative splicing as a mechanism to increase the coding content of genes was emphasized by the accumulation of transcriptomic data over the past decade. In humans about 95% of genes have detectable alternative splice forms (Pan et al., 2008). The nematode is a powerful model to explore networks involved in alternative splicing regulation and the physiological impact of their perturbation (Barberan-Soler et al., 2009, 2011, Zahler, 2012). Recent efforts have been made to systematically identify all possible splice variants of the complete *C. elegans* genome using transcriptome sequencing (RNA-seq) (Ramani et al., 2011, Gerstein et al., 2014, Hillier et al., 2009, Kuroyanagi et al., 2014, Ragle et al., 2015). Most of these analyses reported previously unannotated splice junctions, indicating that saturation has not yet been reached.

The study of individual alternative splicing events can also be performed using fluorescent reporters *in vivo* (Kuroyanagi et al., 2006, 2007, 2013, Ohno et al., 2008, 2012, Tomioka et al., 2016). To date this is the most efficient technique to perform genetic screens in order to identify splicing factors that regulate those events and open the path to study the biochemical details of the regulatory interactions(Amrane et al., 2014, Kuwasako et al., 2014, Mackereth, 2014).

While the power of reporter minigene approaches is undeniable, a major drawback of the method is that it relies on accurate genome annotation of alternative splicing events to build functional reporters. Moreover, current genome annotations make it difficult to estimate the functional relevance of predicted isoforms. As a result, the lack of quantitative annotation can lead to research efforts being unnecessarily spent investigating putative isoforms that are too weakly expressed to be detected *in vivo*.

Here we use the wealth of accumulated RNA-seq data to generate a compendium of quantitative measurements of alternative splicing for each gene in the nematode genome.

## RESULTS

### Characterisation of the spliced RNA-seq compendium

The experiments were selected based on recognition of the keywords “RNA-seq", “transcriptome”, and “*C. elegans* “ and were downloaded from the NCBI Sequence Read Archive (SRA - https://www.ncbi.nlm.nih.gov/sra). We collected RNA-seq data from 96 individual studies and retrieved the raw reads from 1,682 sequencing runs corresponding to 1,185 individual experiments (Full list in Supplementary Table 1). The cumulative number of sequencing reads at our disposal reached a total of 50,544,023,034. Our goal was to measure the relative exon usage for each alternative splicing event, one potential strategy would have been to use exonic sequence coverage, but this method would miss small variations around exon junctions and is complicated by uneven exon coverage. We decided, instead to focus on the 6,631,116,146 reads that spanned a junction between two exons in order to also exclude reads potentially corresponding to contamination by genomic DNA.

We also identified ~287 million reads corresponding to potential trans-splicing of a splice leader sequence to an exon (SL-reads) (Conrad et al., 1991, 1995, Spieth et al., 1993). We found that 97.4% of these trans-splicing events were not detected with good reproducibility (less than 100 reads each) whereas 86% of the SL-reads came from the 36,000 most detected junctions (Figure 1B). These spliced reads were mapped to 667,779 individual splice junctions. We found that ~79% of these splice junctions were not detected with good reproducibility (less than 100 reads each over 1,682 RNA-seq runs). In contrast, 97.6% of the reads came from robustly detected junctions (Figure 1B).

**Figure 1:**
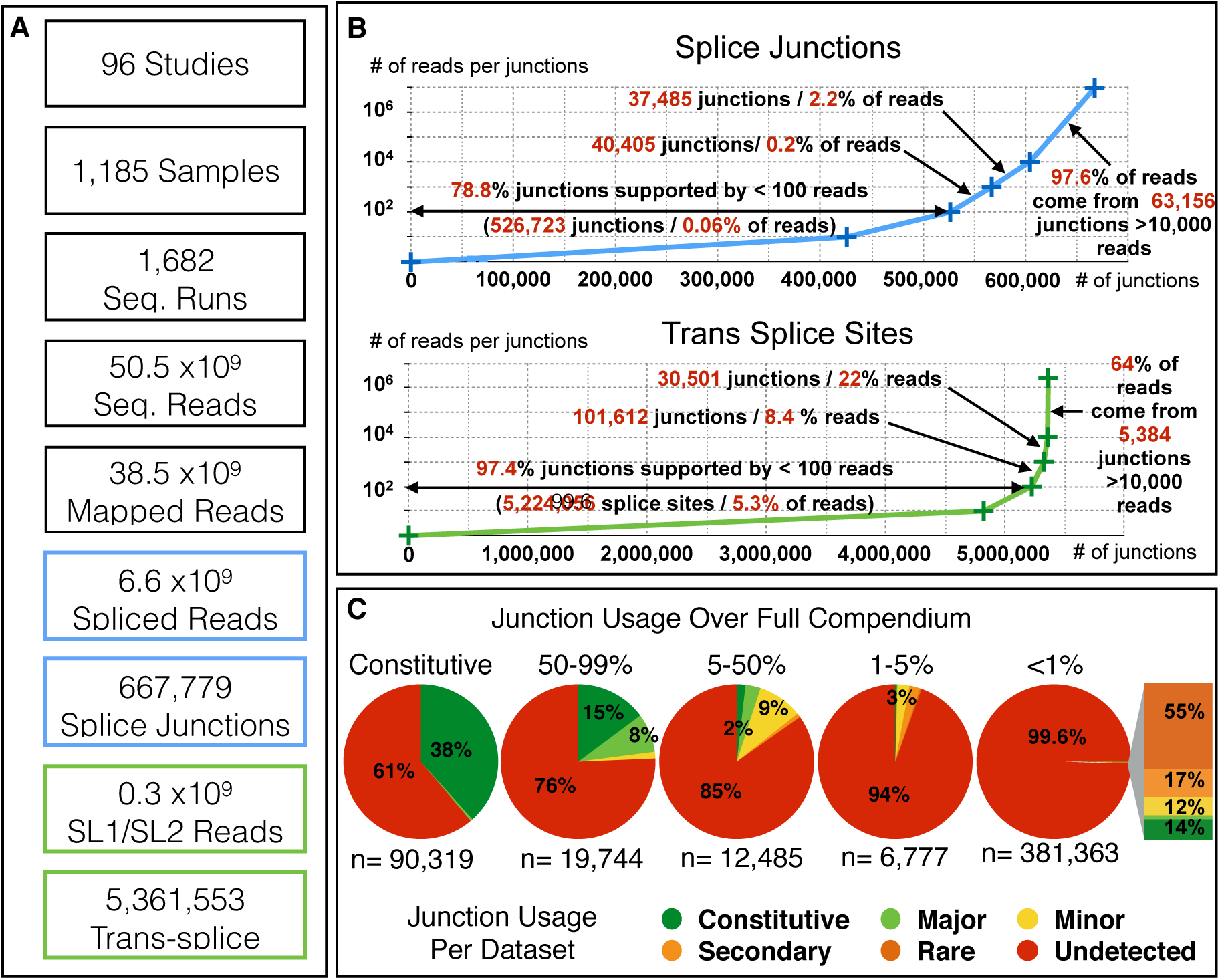
**A)** Relevant figures regarding the compendium dataset used in this study. **B)** Distribution of sequencing reads spanning individual exon/exon (top), or Splice-leader/exon junctions (bottom). **C)** For each splice junction we measured a usage frequency over the whole compendium (see Methods) and assigned them to a usage category (>99%: Constitutive, 50-99%: Major, 5-50%: minor, 1-5%: secondary,, <1% rare). We then evaluated for each junction what their usage frequency was in each of the individual 1,682 RNA-seq runs. The distribution within each category is represented as pie charts. The breakdown of junction with frequency <1% found “non rare” in individual sets is shown as a stacked graph (see also Supplementary Table 2)

Absolute reads threshold are often used in literature for accepting / rejecting events based on a single RNA-seq experiment. Here, however we decided to relate the level of detection for a given isoform to that of other splicing events of the same gene to account for genes with low expression level. We used each splice junction usage frequency to attribute it to the following categories: Constitutive, Major, minor, secondary and rare (in decreasing order of frequency). We checked for the robustness of this classification by comparing it systematically with the usage frequency that would have been obtained using every single RNA-seq run separately (Figure 1C). This analysis confirmed that the Compendium approach provides a more comprehensive evaluation than any single experiment could by improving the dynamic range of detection and averaging out the detection variability between individual sequencing runs. It also shows that the overwhelming majority of the “rare” splicing events are mostly undetected in any given RNA-seq run (see also Supplementary Figure 1).

### Rare isoforms are mostly found in highly expressed genes and are less conserved

While some of the least detected exon junctions come from genes with very low RNA-seq read counts in our compendium - possibly corresponding to genes with restricted spatiotemporal expression - a large fraction come from genes that have been highly covered across a majority of the experiments we analysed. For each alternative splicing event we computed a usage frequency defined by the number of reads found for a given junction divided by the number of reads mapping to any junction sharing an acceptor (or donor) site with it. We found that splice junctions detected 100-fold less than the corresponding main product of their gene (i.e. “rare” variants) represent: 88% of all detected junctions. To investigate whether gene expression level correlates to the amount of detected splice variants, we generated 20 bins of ~1,000 genes grouped according to their expression level in our compendium dataset. We then measured the average number of distinct splicing events per gene in each expression bin. The total number of detected junctions per gene increases with the gene expression level (with the highest expression bin having on average ~70 potential introns), however, if we only count the number of junctions with a usage frequency >1%, we see that the average number of introns per gene is almost invariant (between 5 and 7) regardless of the expression level (Figure 2A). This seems to indicate that these low frequencies junctions do not in fact correspond to functional introns, but rather to accidental misfiring of the spliceosome.

**Figure 2:**
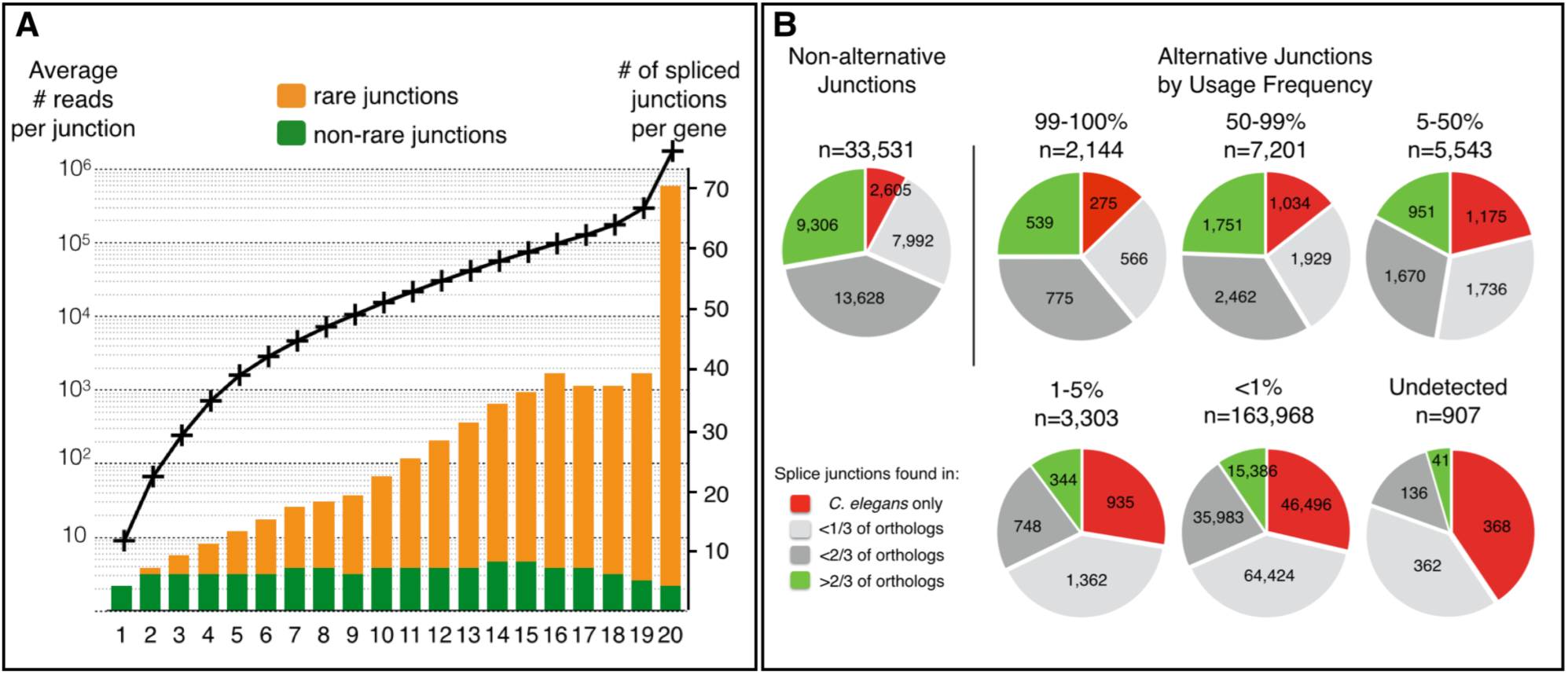
**A)** Introns with low inclusion rate are overrepresented in highly expressed genes. All *C. elegans* genes were ranked according to their expression level (defined by the highest read count for an intronic junction of that gene) and split in 20 bins of ~1,000 genes. For each bin (x-axis) the average expression level (black curve, left axis) and the number of observed splice junctions per gene are plotted. Rare junctions (with an inclusion rate below 1%) are represented in orange and commonly used junctions in green on the bar graph (axis on the right). **B)** Frequently used splice sites are more conserved than rarely used ones. Conservation analysis for pairs of donor and acceptor splice sites grouped by relative inclusion level. For each gene the genomic sequence was aligned with all available orthologs from seven nematode species (See Supplementary Methods). The indicated conservation fraction is the number of genes for which both sites were present, divided by the number of orthologous genes identified.

To evaluate the possible functionality of those rare isoforms we analysed the conservation of all introns boundaries. For each gene with an alternative splice form we retrieved from the WormBase database (http://www.wormbase.org/) the sequence of all available orthologs from seven nematode species (*C. briggsae, C. brenneri*, *C. remanei*, *C. japonica, C. sinica*, *C. angaria*, and *C. tropicalis*). After pairwise alignment of the full gene sequences to each of their counterpart we looked for the presence of paired splice donor and acceptor sites at the expected corresponding positions. As a reference we measured the conservation level for the constitutive intron boundaries within those genes: ~28% (9,306 junctions) were found in at least two-third of the identified orthologs, and only ~8% (2,605) were not found outside *C. elegans.* For alternative junctions used in over 50% of the detected messengers the conservation level is very similar (24-25% found in at least two-third of orthologs and 12-14% specific to *C. elegans*). For junctions with lower level of inclusion we observe progressively less conservation and only ~9% of rare introns are found well conserved while ~29% are not found in other species. We also found that the set of 907 predicted junctions (from Wormbase) for which no RNA-seq evidence could be found in our compendium shows even less conservation, as could be expected for non-functional sequences (Figure 2B).Overall, we find that “rare” junctions are less evolutionarily conserved than more frequently used junctions.

The number of reads per junction ranges over six orders of magnitude depending of the gene. This means that 1,000 reads can be the maximum count for a gene while representing only 0.01% of reads for another, we therefore propose that the usage frequency is a better predictor of the functionality of a splice junction than raw read count.

### Trans-splicing is potentially ubiquitous in *C. elegans*

In *C. elegans* mRNAs the 5’ UTR is often cleaved off and replaced by a splice leader (SL) sequence that is provided by an independently transcribed snRNA (Conrad et al., 1991, 1995, Spieth et al., 1993). Splice Leader 1 (SL1) is mostly associated with genes linked to a proximal promoter directly upstream while SL2 is generally associated with genes located downstream in polycistronic operons. Below, we therefore refer to trans-splice sites as indicative of a “gene start” rather than transcription start.

We detected trans-spliced sites for most protein coding genes but found that a vast number of trans-splice sites with very low read counts likely correspond to biological noise. This is supported by the observation that highly expressed genes have larger number of trans-splice sites while the average number of robust splice site per gene is independent of the expression level (Figure 3A). As for cis-splicing, the wide variation of gene expression levels precludes using an absolute read-count threshold to identify *bona fide* trans-splicing sites. Therefore, to filter out potential spurious trans-splicing events, we decided to consider “robust” the sites that were corroborated by at least 25% of the SL-containing reads associated to that gene. While this definition is not perfect, it provides a good approximation of the main SL-sites used for most genes. We found ~20,000 robust trans-splice sites spanning 17,060 genes (84% of protein coding genes). Previous work by Allen *et al*. attempted to systematically identify all trans-splicing sites in the *C. elegans* genome. They detected 70% of genes subjected to trans-splicing and found a tendency for highly expressed genes to be more trans-spliced than the less expressed genes (Allen et al., 2011).

**Figure 3:**
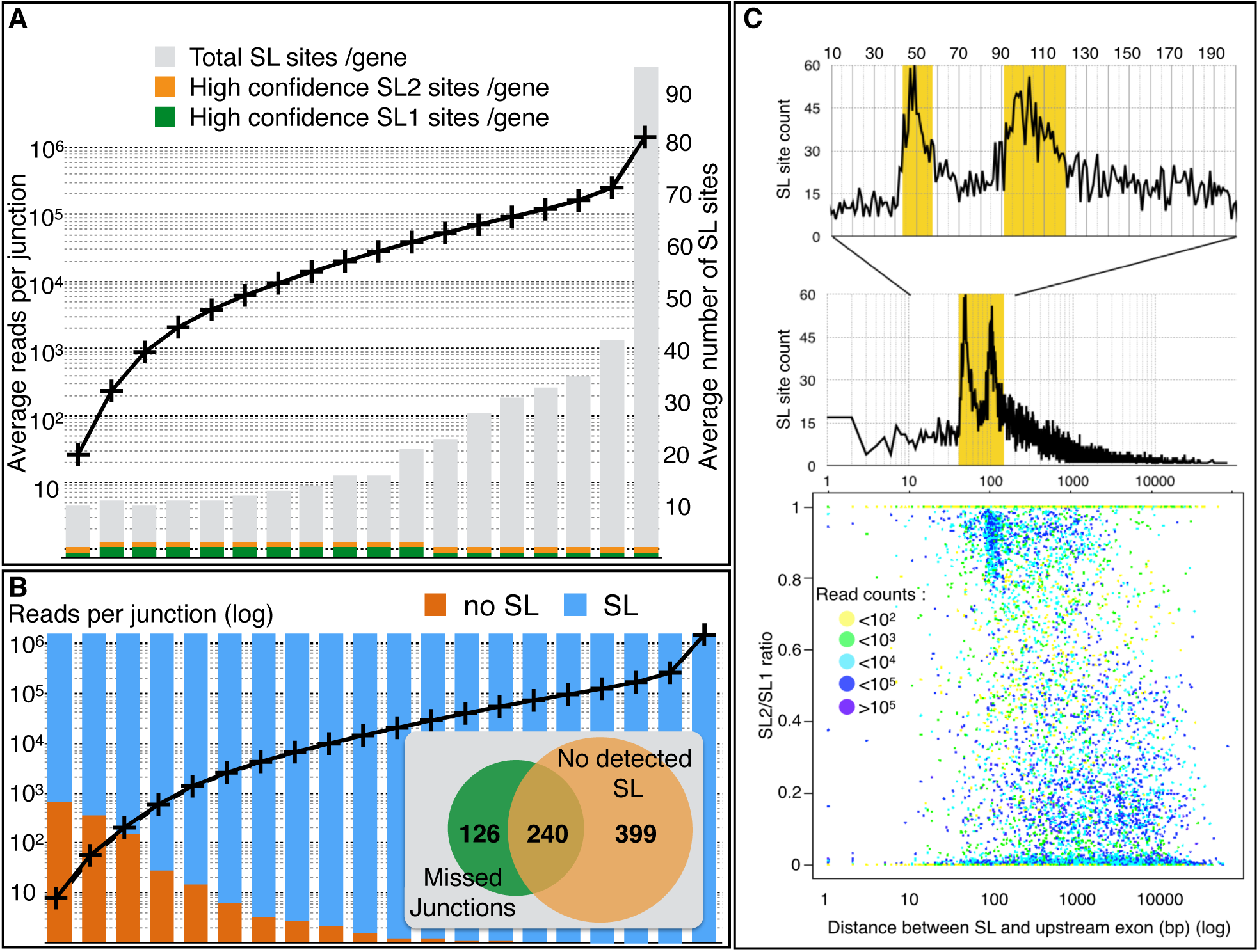
**A)** All genes for which at least one robust SL site was detected were ranked according to their expression level (as defined highest read count for a intronic junction of that gene) then grouped in bins of ~1000 genes. For each bin the average expression level (left axis) and the average number of SL site per gene found are plotted. Here we counted as high confidence sites loci that contained at least 25% of the SL-containing reads of that gene. **B)** Genes without detectable trans-splicing have mostly low expression levels. All *C. elegans* genes were ranked according to their expression level (defined by the highest read count for an intronic junction of that gene) and split in 20 bins of ~1,000 genes. For each bin the average expression level (black curve, left axis) is plotted. For each bin we represented the fraction of genes for which no SL junction was found (orange) and for which at least one robust SL site was found (blue). For the 1,000 genes with the lowest expression level we inserted a Venn diagram representing the number of genes for which either some predicted cis-junctions were missed and/or no SL site was detected. **C)** A scatter-plot of the ratio of SL2/SL1 found at each robust splice site against the distance to the closest upstream exon shows a discernible bias for sites with over 80% SL2 splicing to be located at ~100 nt downstream of the nearest exon. Each dot is colored according to the number of sequencing reads supporting the corresponding SL site. A zoomed plot of the number of occurrences of SL sites at each distance shows two sharp peaks indicative of a strong preference for SL2 splicing to occur either at a distance of ~50 nt or ~100 nt downstream of the previous exon.

In the 2011 study, RNA-seq reads were mapped against a database of potential trans-splice sites generated *in silico* by joining all SL sequences with all annotated acceptor splice sites. In contrast, our detection method is independent of the accuracy of the genome annotation and therefore allowed us to recover trans-splice sites that had been previously missed. In addition to the difference in mapping strategy we also benefited from using a 1000-fold larger number of reads. Most of the 15% of genes for which we found no SL-site are among the lowest expressed genes in the genome (Figure 3B). This suggests that the RNA-seq coverage for these genes hasn’t been saturated even by our compendium of datasets and that more targeted approaches directed at these low expression genes are needed to determine their trans-splicing status (Supplementary Figure 2.

For each of the robust SL-sites we determined what proportion of SL1 or SL2 splice leader sequence was used at each position (expressed as SL1/SL2 ratio). We then compared this ratio to the distance to the closest upstream exon (Figure 3C). We found that most positions are strongly favoured by one SL sequence rather than the other and that SL2 trans-splicing is strongly preferred when the closest exon is located either ~50 or ~100 nucleotides upstream of the acceptor site, thus confirming and refining the previously reported observations (Allen et al., 2011).

### Genome-Wide Quantitative Visualization of Differential Exon Usage

For each gene we generated a visual summary of the splicing data we collected (~20,000 images collated as Supplementary Figure 3). For each gene we represented only the most commonly used exons (i.e.:constitutive exons and the most common forms for alternative splicing events) and the quantitative values of all the spliced junctions obtained from our analysis. This allows to see at a glance what is the main product of any given gene, and what fraction of the messengers use alternative forms. In addition, we also represented the identified trans-splicing positions and the level at which they were detected. This allows for the first time to directly see how many distinct promoters are used by a given gene and their relative contribution to that gene’s expression.

To illustrate the usefulness of our visual representations, we present in Figure 4 three examples together with their current genome browser representation in WormBase, including all annotated isoforms. For the first two examples, alternative isoform expression has been well described and validated by fluorescent reporter constructs:

- DAF-2 protein is *C. elegans* insulin/IGF receptor ortholog well known for its effect on lifespan extension and *Dauer* formation (Kenyon et al., 1993; Apfeld and Kenyon, 1998). Immunodetection of the protein indicates that it is predominantly expressed in a subset of neuronal cells in the animal nerve ring (Kimura et al., 2011). It has recently been shown that an alternative functional isoform generated by inclusion of a cassette exon (exon 11.5) is expressed more specifically in sensory neurons and particularly in starvation conditions (Ohno et al., 2014, Tomioka et al., 2016). We found that this isoform accounts for ~15% of the detected reads (~5,000 reads exon 11.5 inclusion vs ~36,000 for skipping, Figure 4A). While we find support for several of the annotated alternative start sites only one major trans-spliced site stands out as the likely main start site for this gene.
- PTB-1 is a hnRNP family protein member whose tissue specific expression is necessary for the inclusion of exon exon 11.5 in *daf-2* (Tomioka et al., 2016). Tomioka and colleagues demonstrated that *ptb-1* was expressed from two alternative promoters active in distinct neuron subsets. Consistent with this observation we found ~15,000 reads for the PTB-1a- or PTB-1b specific splice junctions and ~30,000 reads for the junctions shared by both isoforms. Our data also support the existence of a third unverified promoter with a lower activity (black arrow). We can also observe that a previously unreported skipping of exon 11 accounts for 16% of the detected junctions (5,439 reads joining exons 10 −12 vs 28,758 reads joining exons 10-11, Figure 4B). This exon skipping causes a frame shift that could generate a truncated version of the protein lacking the last 167 amino-acids.

**Figure 4:**
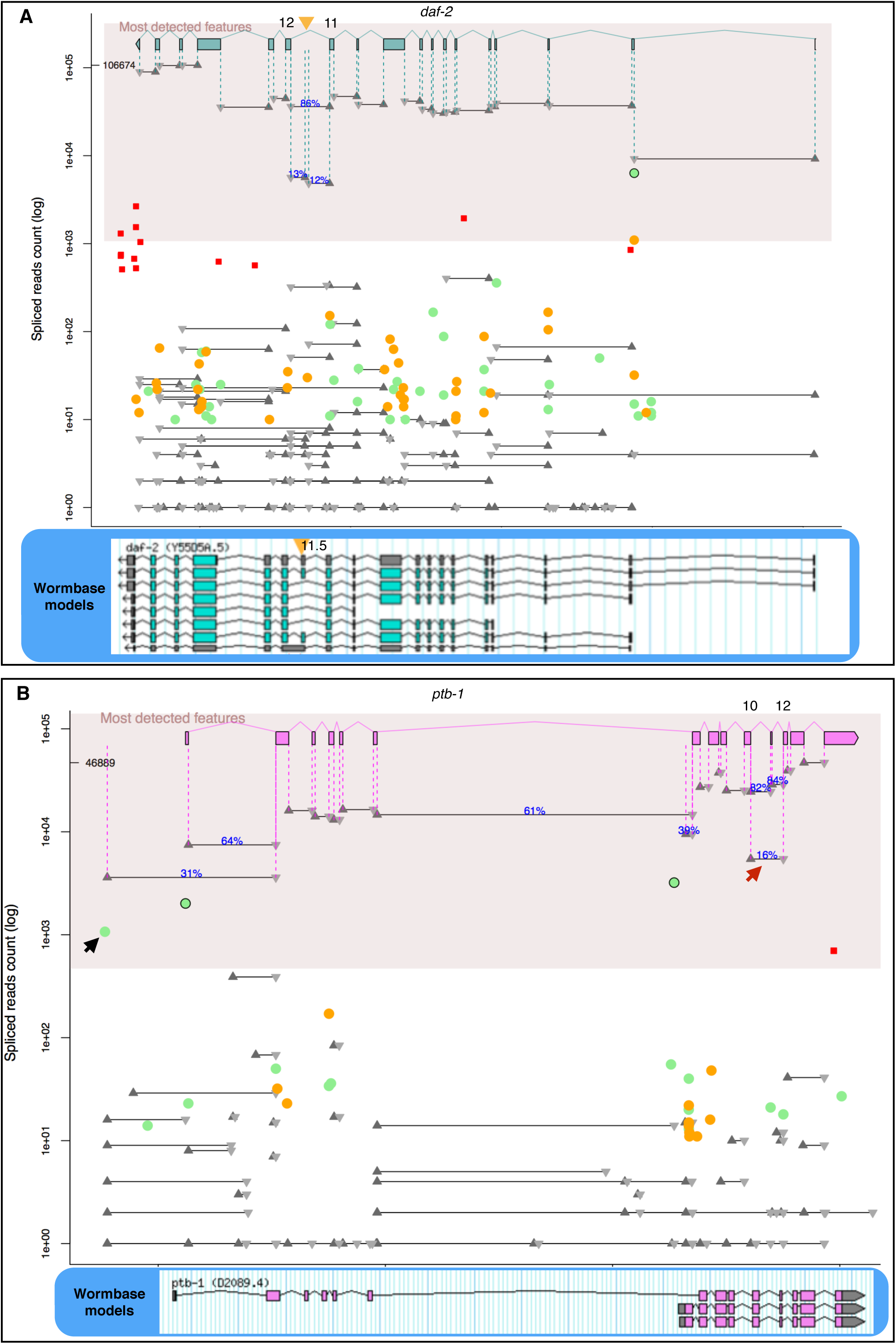
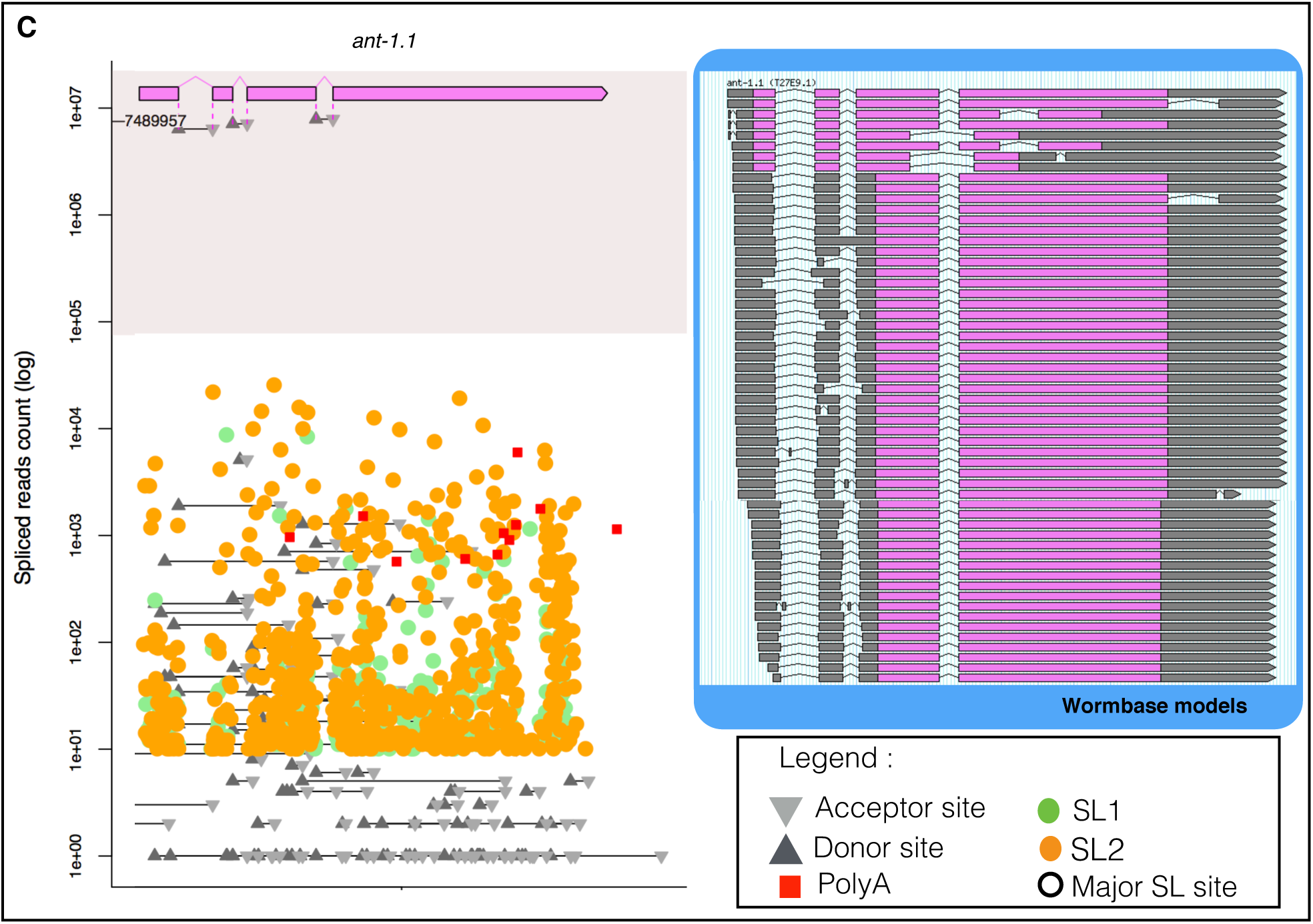
Quantitative visualisation of relative splice-sites usage. We present a gene model constituted of the most commonly detected exons, associated with the absolute read count for each cis- and trans-splicing event in the gene, as well as detected polyA tail addition, on a logarithmic scale. We highlighted the area containing high confidence events (detected in at least 1% of the transcripts) and included junction usage frequencies for alternative events. As a reference we include the current Wormbase model **A)** *daf-2*: exon junctions corresponding to the inclusion of exon 11.5 (orange arrowhead) specific to a small subset of neurons during starvation are detected at ~15% of the level of other junctions. Evidence of trans-splicing at exon 2 and a 10-fold increase in read numbers between introns 1 and 2 seem to indicate that this is in fact the most used expression start for this gene **B)** The *ptb-1* gene has two confirmed alternative promoters active in two distinct neuronal subsets. In our data analysis this manifests as two Major SL1 trans-splicing junctions associated to the detection of the corresponding isoform-specific exon-exon junctions. Note how common junctions are detected with a level corresponding to the sum of both isoform-specific rates downstream of the second promoter. Our data seem to support the existence of a third unverified promoter with a lower activity (black arrow). We also detect an exon junction between exon 10 and exon 12 indicating that about 10% of the transcripts are skipping exon 11 (red arrow). **C)** The *ant-1.1* gene had over 50 isoforms predicted in Wormbase. Our RNA-seq analysis detected the three constitutive junctions with over 6 millions reads each. All other junctions are orders of magnitude below the overall expression level of this gene indicating that there is only one functional isoform of ANT-1.1.

From these (and several other) examples we concluded that our visualisation tool was indeed suitable to provide a measurement of isoform usage that is consistent with known alternative splicing events even when they are pertaining to a very restricted number of cells.

We next wanted to see if we could as easily interpret our data for genes for which no prior information on alternative splicing patterns was available. As an extreme example we show here *ant-1.1* which had over 50 predicted isoforms in the *C. elegans* genome annotation WS251. ANT-1.1 is an essential mitochondrial adenine transporter that is ubiquitously expressed (Farina et al., 2008). Our data clearly discriminates between two classes of exon-exon junctions for *ant-1.1*: on one hand three constitutive junctions for each of which ~6 million reads are found, and on the other hand 170 junctions supported by a number of reads ranging from 1 to 200,000. This second class of junctions, amounting to a very small proportion of the detected RNA products (2 to 6 orders of magnitude less than the main product), is likely to result from stochastic misfiring of the splicing machinery as most of the isoforms produced contain frame-shifts and premature stop codons (Figure 4C). We considered these splice forms as “rare variants” in our curation.

## DISCUSSION

In eukaryotes, the accumulation of aberrantly spliced messengers that could encode potentially deleterious truncated proteins is prevented by Nonsense Mediated Decay (NMD) (Jaillon et al., 2008, Farlow et al., 2010). Species devoid of NMD tend to almost entirely lack introns (Lynch, 2006) indicating that an error proof splicing system has not arisen through natural evolution (or is yet to be discovered). Together, these observations support the widespread existence of biological noise in the splicing process.The sensitivity of modern deep sequencing methods for transcriptome characterisation ensures that even rare aberrantly spliced messengers will be detected alongside functional splice forms. Traces of messengers targeted for decay, that are accumulating in NMD mutants, can be detected in wild type individuals as well (Barberan-Soler et al., 2009, Ramani et al., 2009). Moreover, a recent study unveiled a significant decrease in splicing accuracy correlated with age in *C. elegans* (Heintz et al., 2016).

We reasoned that by exploring a large collection of RNA-seq datasets we could compile a nearly comprehensive list of splice junctions in the *C. elegans* genome. The total amount of data used in our study provides a robust quantitative measure of the frequency of usage of each detected alternative splice junction and the expanded dynamic range obtained also allows for discrimination between genuine alternative splicing and potential biological noise. Our observation that “rare” junctions come disproportionately from highly expressed genes and are less conserved in other nematode species, could indicate that most of these junctions correspond to biological noise causing accidental splicing outside of the preferred functional sites. A similar analysis in human cell lines also found that rare alternative splicing events are more frequently observed in highly expressed genes and tend to be less conserved across species (Pickrell et al., 2010). The authors of that study also found that the small fraction of reads coming from unannotated rare junctions covered a large number of likely spurious splicing events.

While it is likely that our discrimination between rare and robust splice variants offers a good approximation for the functionality threshold of any given isoform, it is almost certain that some exceptions will apply. It is possible, for example, that a ubiquitously expressed gene also encodes a rare variant with a limited cell specificity, but studying and validating this kind of event will constitute its own challenge. In that context our classification should be considered a warning sign: these “rare” events are unlikely to be functional and studying them will be not be trivial as they are not detected in most RNA-seq experiments.

Our compendium based meta analysis provides a widely expanded dynamic range of detection with genes having between 0 and 10^7^ reads per junctions. If we considered every splice junction detected by any RNA-seq experiment in our compendium we would conclude that ~94% of *C. elegans* genes are submitted to alternative splicing (Table 1). If we consider only genes for which there is a second isoform with a frequency of at least 1% of the major isoform, this number drops to 35% (~7,000 genes). This could be a valid definition since our analysis suggests that the majority of “rare” splicing events corresponds to biological noise rather than a conserved functional mechanism. However, it is possible that for some genes a rare isoform indeed is critical for a cell-specific function. Conversely, we cannot proclaim that every event that is above the 1% threshold is a genuine alternative splicing event. If we place the bar at 5% of the gene expression level, then only 4,700 genes have more than one isoform. There is no objective quantitative criteria that can systematically discriminate between functional and spurious alternative isoforms at this time.

**Table 1:**
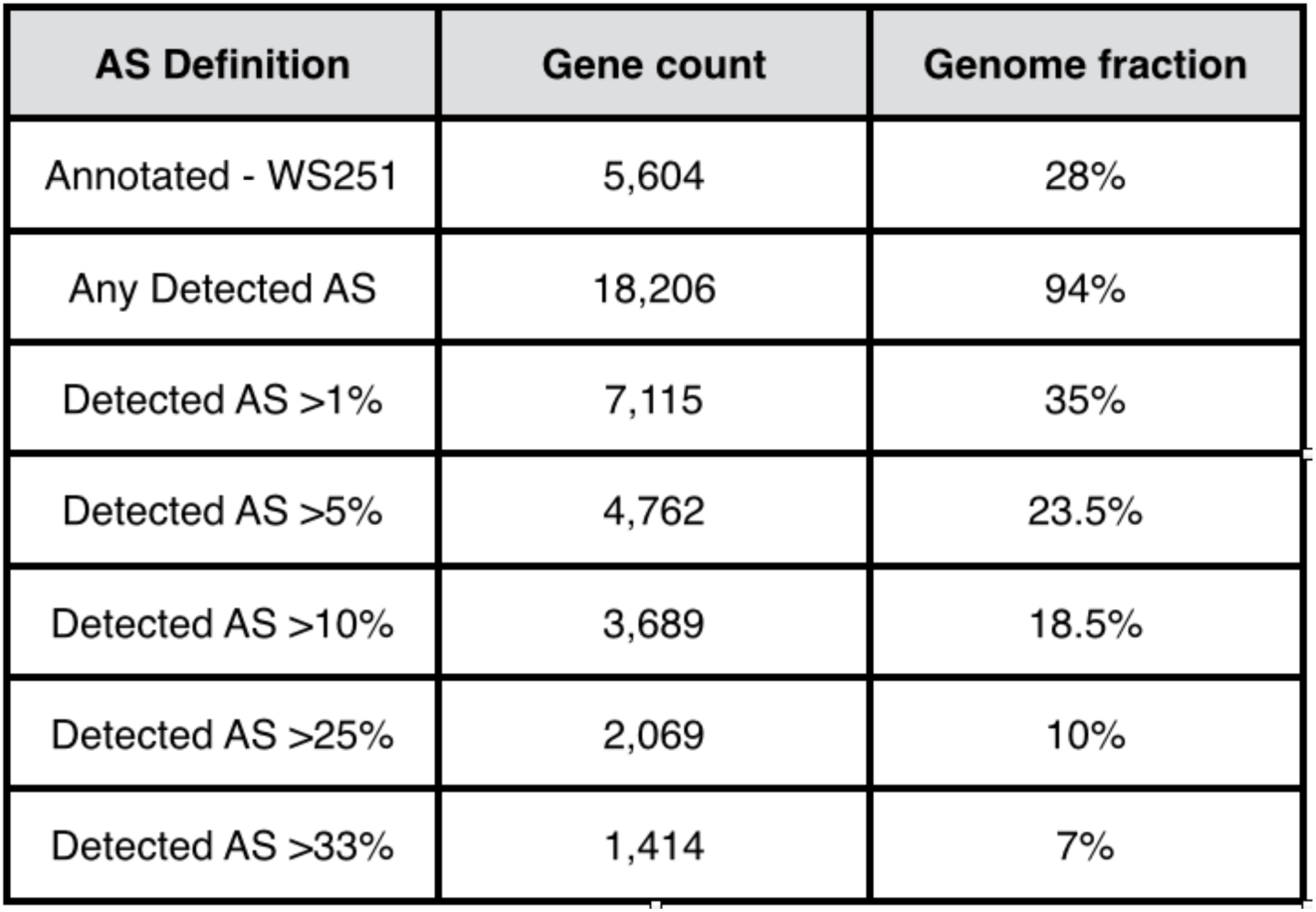
Evaluation of the proportion alternatively spliced (AS) genes in the *C. elegans* genome. The threshold used to define genuine alternative splicing has a major influence on the estimation of the prevalence of the phenomenon.

The visual representation we propose here allows to immediately see if a gene has a set of genuine introns clearly separated from background splicing noise, or if there are intermediate splice variants worthy of investigation. For genes with multiple alternative promoters our representation also provides a visual ranking of the relative strength of each promoter which can be indicative of their tissue specify. Such first approximations will be very useful for generating hypotheses and designing experiments to test them. Importantly these representations are unbiased and objectively derived from a wide range of independent experimental observations.

The combination of increased depth of sequencing data with an unbiased SL site detection strategy led us to find evidence of trans-splicing for 84% of *C. elegans* protein coding genes (versus 70% previously). We were not able to detect SL sites mainly for genes that have the lowest expression level. This indicates that despite the large dynamic range of our data compendium we didn’t reach sufficient coverage for those genes. Since even the accumulation of several hundred full-animal RNA-seq data collection did not provide the sensitivity needed, more targeted RNA-seq experiments will be necessary to explore the function and specificity of the least expressed genes and their isoforms (Fox et al., 2005, Hashimshony et al., 2012, Schwarz et al., 2012).

Current RNA-seq-based estimations claim that greater than 95% of human multi-exon genes express multiple splice isoforms (Pan et al., 2008, Wang et al., 2014) which is very similar to the number we find in *C. elegans* when considering every detectable isoform. The validity of this evaluation is contested by some other transcriptomic and proteomic studies (Pickrell et al., 2010, Tress et al., 2017). Applying our strategy of aggregating large number of RNA-seq datasets to flag potential stochastic splicing could help shed new light on the question of the prevalence of alternative isoforms and proteome complexity in other metazoan organisms.

## METHODS

### Datasets selection

Sample experiment and run accession numbers and sequence files were downloaded programmatically using the NCBI “E-utilities” tools “esearch” and “efetch” (https://www.ncbi.nlm.nih.gov/books/NBK25501/) and the Unix utility “wget”, and read files in FASTQ format were generated using the “fastq-dump” tool from the NCBI SRA Toolkit 2.5.0. Illumina mRNA-seq experiments were selected with a regular expression search of the experiment descriptions, leading to a final compendium of 1,682 datasets (1,214 single-end and 468 paired-end). Read files were taken as is; no read processing was performed.

### Exon-exon junction identification

Junctions containing canonical intron splice sites (GT-AG, GC-AG, and AT-AC) were identified by mapping raw RNA-seq reads to the *C. elegans* genome sequence (obtained from WormBase release WS251, http://www.wormbase.org/) using TopHat2 2.0.14 (Kim et al. 2013) based on the Bowtie2 2.2.5 (Langmead and Salzberg, 2012) core read aligner. The list of junctions and the number of reads spanning them were retrieved from the “junctions.bed” output file of TopHat2. All datasets were mapped in single-end mode and the following options were set for TopHat2 (all other parameters as default): “--min-intron-length 10 --max-intron-length 20000 --read-mismatches 3 -- read-gap-length 2 --read-edit-dist 3 --max-multihits 2 --b2-sensitive --segment-mismatches 2 --segment-length 15 --min-segment-intron 10 --max-segment-intron 20000 --no-coverage-search”. RNA-seq reads having only a few matches to one exon side of a junction may be missed by the above mapping strategy. To find additional junctions and additional reads corresponding to the previously detected junctions, unmapped reads were re-mapped to the *C. elegans* WS251 genome by setting the following TopHat2 parameters to force the detection of deletions of up to 1 kb in length: “--read-gap-length 1000 --read-edit-dist 1003 --b2-ma 3 --b2-rdg 3,1”. Sequences of 200 bp flanking each side of a given deletion site were joined and then mapped to the *C. elegans* genome with TopHat2 run with the same parameters as in the original search to recover further reads containing canonical exon-exon junctions.

In addition, junctions that were predicted in WormBase WS251 but that were not recovered in our searches (“undetected junctions”) were included. We excluded junctions corresponding to putative introns larger than 2kb from the downstream analyses.

### Junction usage quantification

The relative usage of splice junctions was estimated as follows: First, all junctions for which at least one boundary was within the coordinates of a gene predicted on the same DNA strand in WormBase WS251 were assigned to that gene. Then, for a given gene, a donor ratio was computed for all detected junctions that shared a common donor boundary by dividing the number of RNA-seq reads mapping to a junction by the sum of reads mapping to the set of junctions having the same donor site. An acceptor ratio was similarly calculated for detected junctions sharing a common acceptor site. If both ratios could be computed, then the usage ratio was set to the ratio that was based on the largest number of reads. In the case of junctions that did not share any boundary with any other junctions in the gene, the usage ratio was set to 1 (constitutive junction). We also defined a ratio relative to the junction with the highest number of mapped reads for the gene (“max_junction”), i.e., number of reads for the junction divided by number of reads for max_junction. If max_junction had no reads (in the case of a totally undetected gene), then the usage ratio was set to “NA” (not available). If a junction “Max_Ratio” was below 1% the junction was classified as “rare”. Note that a junction may be assigned to more than one gene (e.g., when genes are overlapping or in close proximity), and will have usage ratios specific to each gene. Supplementary Table 2 contains, the genomic positions, the number of supporting reads in our compendium, the number of experiments that detected it, the inclusion ratio and the curated category for all exon-exon junctions we detected.

### Conservation analysis

A comparative analysis was conducted to determine whether exon-exon junctions identified in *C. elegans* were conserved in other nematodes. The list of genes that were predicted to be orthologous between *C. elegans* and seven other *Caenorhabditis* species (*C. angaria*, *C. brenneri*, *C. briggsae*, *C. japonica*, *C. remanei*, *C. sinica*, and *C. tropicalis*) was retrieved from WormBase WS251, as well as the corresponding gene annotation and genome sequence files. In a given species there may be several predicted orthologs to the same *C. elegans* gene. A global and optimal pairwise alignment was computed according to the Needleman and Wunsch algorithm {Needleman and Wunsch, 1970) between the full nucleotide sequence of each *C. elegans* gene and its ortholog(s) using the program “needle” from the EMBOSS 6.6.0 package (Rice et al. 2000). Then, alignments were scanned for junctions. For each *C. elegans* exon-exon junction along the gene, if the four bases at the corresponding alignment positions in the other species matched a canonical splice site (GT-AG, GC-AG, or AT-AC), then the junction was considered as conserved between *C. elegans* and the other species (the junction need not to be identical between the two species to be considered present). Supplementary Table 3 contains the data pertaining to this conservation analysis,

### Trans-splice site identification

Trans-splice sites were identified from the RNA-seq reads that did not map to the *C. elegans* genome (with or without introns). Our strategy was multi-step (See Supplementary Figure 4). First, cutadapt 1.3 (Martin, 2010) was employed to identify all sequencing reads containing a putative SL sequence (or the 3’ end of it) and to extract the sequence downstream of the SL. The search requirements were: a match length of at least 5 nt to the SL with a max. of 10% mismatches and a downstream sequence of at least 15 nt (options “-e 0.10 -O 5 -m 15 --trimmed-only”).

The following SL1 sequence "CTCAAACTTGGGTAATTAAACCG" and seven SL2 variant sequences ("GGTTTAAAACCCAGTTACCAAGG", "GGTTTTAACCCAGTTAACCAAGG", "GGTTTTAACCCAGTTACTCAAGG", "GGTTTTAACCCAGTTTAACCAAGG", "GGTTTTAACCCATATAACCAAGG", "GGTTTATACCCAGTTAACCAAGG", and "GGTTTTAACCCAGTTAATTGAGG"), and their reverse-complement, were used as queries for the search. Then, three mapping algorithms were used in succession to determine the genomic locations of the corresponding *trans*-splice sites. The read portion downstream of the SL portion was first mapped to the *C. elegans* WS251 genome with TopHat2 (with options “--min-intron-length 10 --max-intron-length 20000 --read-mismatches 3 --read-gap-length 2 --read-edit-dist 3 --max-multihits 2 --b2-sensitive --segment-mismatches 2 --segment-length 15 --min-segment-intron 10 -- max-segment-intron 20000 --no-coverage-search”), which allows reads to contain introns but requires full alignment. Then, the unmapped SL reads were mapped to *C. elegans* WS251 with Bowtie2 - which does not make spliced alignments but allows for partial mapping with soft-clipping. Bowtie2 was run with options “--local --sensitive-local” (all other parameters set as default). Finally, the remaining unmapped SL reads were aligned to the *C. elegans* genome with GSNAP from the GMAP-GSNAP package release 2017-01-14 (Wu and Nacu, 2010), which allows for both introns and soft-clipping. The following GSNAP options were set (all other parameters as default): “-- nofails --novelsplicing=1 --localsplicedist=20000 --novelend-splicedist=20000 --suboptimal-levels=0 --max-mismatches=3 --indel-penalty=2 --max-middle-insertions=2 --max-middle-deletions=2 --max-end-insertions=2 --max-end-deletions=2 --input-buffer-size=100000 --output-buffer-size=100000”. In the above mapping strategy, in case of soft-clipping the genomic position of the *trans*-splice site was shifted according to the length of the clipped region. Otherwise, it was taken as the mapping position of the 5’ end of the SL read (i.e., the 5’ end of the sequence downstream of the SL).

To further improve the quantification of the number of reads spanning a given SL position, full reads that mapped to the *C. elegans* genome and that extended a few bases over the discovered SL positions were examined. For those reads, the sequence portion extending beyond the SL position was compared with the corresponding genomic sequence at that position and with the 3’ ends of the SL1 and SL2 variants. If the number of mismatches to any of the SL ends was smaller than that to the genomic region, then the read was considered as having a SL piece and was added to the count of reads spanning the given SL position.

Intergenic SL sites were assigned to the nearest downstream gene if it was located within a distance of two kb. Supplementary Table 4 contains the data pertaining to the trans-splicing events with a read count >10.

### Non-genomic polyA site identification

From the set of RNA-seq reads that did not map to the *C. elegans* genome, cutadapt was used to identify those that harbored a polyA stretch at their 3’ end and extract the upstream read region. Reads carrying a stretch of at least 10 A residues, with a max. of 20% mismatches, and a remaining upstream portion of at least 15 nt were searched for (cutadapt options “-e 0.20 -O 10 -m 15 --trimmed-only”). The upstream portion of these reads was then mapped to the *C. elegans* genome with TopHat2 (run with the same parameters as for the *trans*-splice site search) and the polyA site location was set to the 3’ end of the read mapping position. Then, sites corresponding to genome-encoded polyA runs were filtered out. For that, genomic polyA’s were identified by searching the *C. elegans* genome sequence by means of the program “fuzznuc” from the EMBOSS package with the query “AAAAAAAAAA” and allowing for two mismatches (options “-pmismatch 2 -pattern AAAAAAAAAA -complement”). Potential PolyA sites whose coordinates matched the genomic sites were discarded.

Intergenic polyA sites were assigned to the nearest upstream gene if it was located within a distance of two kb. Supplementary Table 5 contains the genomic location for polyA sites with a read count >500.

### Graphical representation of quantitative splicing and *trans*-splicing

For each gene a graphical representation showing the gene features (exons, splice junctions, SL and polyA sites) along with quantitative usage data was generated using R 3.2.0 R Core Team 2015 (http://www.R-project.org/). We present a gene model constituted of the most commonly detected exons. On a logarithmic scale we report the absolute read count for each cis- and trans-splicing events, and polyA additions (the read count for the most detected cis-junction is indicated on the y-axis). Vertical dashed lines connect the non-rare junctions to the gene model. Usage ratio of alternative events is indicated. We highlighted the area containing features detected at a level of at least 1% of the maximum junction read count for the gene (shaded area). All plots are shown in Supplementary Figure 3.

## DATA ACCESS

The raw data used in this study are publicly available and were taken for the NCBI SRA repository. Graphical summaries of our quantitative spicing analysis for each individual gene is accessible as Supplementary Figure 2 and will be made available on Wormbase.

## ACKNOWLEDGMENTS

This work has been funded by Inserm (DD, NJT) and the French Ministry for Higher Education and Research (JRMM). We thank Dr. Axel Innis for providing access to his computer cluster at the IECB, and the University of Bordeaux for access to the supercomputer of the Mésocentre de Calcul Intensif Aquitain (MCIA).

